# Long term hybrid zone dynamics in red- and yellow-bellied toads estimated from environmental data at allopatric and parapatric scales

**DOI:** 10.1101/2025.04.24.650419

**Authors:** Jan W. Arntzen

## Abstract

**Aim:** Aims of the study are to identify environmental parameters underlying the mutual distribution in a species pair engaged in a long and winding hybrid zone, to reconstruct pattern and process of species range developments following the Last Glacial Maximum (LGM), and to open research lines for the study of hybrid zone dynamics in a model system.

**Location:** Central and Eastern Europe, Croatia.

**Taxon:** The red-bellied toad *Bombina bombina* and the yellow-bellied toad *Bombina variegata*.

**Methods:** The construction of two-species distribution models (TSDM) at allopatric and parapatric scales and the reconstruction of geographical clines across a low mountain range, in transects with and without extensive lowland forest cover.

**Results:** It is confirmed that *B. variegata* is a mountain species and *B. bombina* is a lowland dweller, with hybrid populations in between, but traditional distribution models are oversimplistic. Environmental parameters selected with TSDM at both allopatric and parapatric scales are elevation and forestation. At transects with lowland forestation hybrid zones are positioned further away from elevated areas than in the absence of lowland forest. *Bombina variegata* stronghold areas are characterized not just by elevation but also by forestation. Out of 17 historical *Bombina* studies that are evaluated for promise in the study of hybrid zone dynamics several qualify for extended research.

**Conclusions:** The overall species mosaic with isolated mountain strongholds suggests that *B. variegata* was displaced from the surrounding lowlands upon its counterparts northward advance, following post-LGM climate amelioration. Initial hybrid zone formation will have been along the lower Danube River, distant from present-day positions. Because lowland forestation constitutes a buffering effect to species replacement, current hybrid zones may not have reached equilibrium conditions, depending on deforestation history and other characters of the landscape. Future genetic work on *B. variegata* enclaves may shed light on the pattern and process of hybrid zone movement and species replacement, though timing and the size and environmental signature of the enclaves will affect genetic introgression in ways that may be hard to disentangle.

## Introduction

Species distribution modelling (SDM) aims to identify the ecological factors that limit and define species ranges and can be used across temporal windows, to reconstruct past distributions and to evaluate the impact that environmental change might have on species extinction probabilities. Therewith, SDM has become a popular tool in fields as far apart as phylogeography and wildlife management (Franklin, 2010; Peterson *et al*., 2011; Guisan *et al*., 2017). The most notable obstacles to meaningful SDM remain identification error, spatial biases in presence data, the inferred nature of absence data and the selection of explanatory variables (Lobo et al., 2010; Soley-Guardia et al. 2024).

Species distribution models are frequently constructed for single species without reference to biotic interactions, although it has been noted that such disregard might give rise to misleading results (Davis *et al*., 1998; Leathwick, 2002; Araujo and Luoto, 2007; Meier *et al*., 2010; Wisz *et al*., 2013). Biotic interactions can be exploitative or mutualistic with positive effect or they can be negative through disease, predation and competition, therewith enhancing or limiting population processes and, ultimately, species distributions. The main reasons for ignoring biotic factors in species distribution modelling are that they may be difficult to parameterize and that blanket coverage is mostly unavailable. For species that heavily interact to the extent that their ranges are mutually exclusive, SDMs may be of limited value, in the same way that the edge of a continent is not helpful in understanding the ecological limitations of a terrestrial species (e.g., Dufresnes et al., 2020ab). In such cases of parapatry the drawback, however, may be turned into an advantage, because species with abutting ranges offer the opportunity to *contrast* their ecological preferences such as in ‘two-species distribution modelling’ (TSDM), in which presence data from two species are compared against the background of environmental data, to yield insights into species habitat differentiation. Parapatry is common, especially among related organisms with low dispersal capability (Key, 1981; Bull, 1991). With the advent of molecular genetic data in taxonomy, mosaics of closely related species continue to be resolved (Marzahn *et al*., 2016; Pyron *et al*., 2022; Yang *et al*., 2022), underlining the ubiquity of parapatric contact zones and, therewith, the scope for TSDMs (for a review in European amphibians and reptiles see Arntzen, 2023a).

Another dimension to parapatry is the occurrence of narrow hybrid zones at the species interface. Hybridization offers insight into speciation and the forces that maintain barriers to reproduction (Ravinet et al., 2017; Peñalba et al., 2024). Hybrid zones provide excellent opportunities to test how the environment shapes barriers to reproduction and hybrid fitness, and how differences in reproductive barriers between two species influence their relative success across habitats. I here apply TSDM and hybrid zone cline analyses to a species pair of fire-bellied toads (genus *Bombina*) that represents a model system in speciation research. Aims of the study are to identify environmental parameters that determine the mutual distribution of *B. bombina* and *B. variegata* that engage in a long and winding hybrid zone across central Europe, and to reconstruct pattern and process of species’ range developments following the Last Glacial Maximum (LGM). It will be argued that, by ignoring important components of the *B. variegata* environment, existing species distribution models are naïve and that a process of hybrid zone movement is ongoing, depending on characteristics of the landscape, therewith opening research lines for the study of hybrid zone dynamics in a model system.

## Material and methods

### Research areas and species distribution data

Areas for research are central Europe that here covers the sector of *Bombina* species range overlap bounded by 14-28 eastern longitude and 43-51 northern latitude (Figure 1), and central Croatia with the sections Jastrobarsko and Pešćenica at opposite flanks of the Vukomerić hills south of Zagreb (Figure 2). Species distribution data are publicly available and used to *construct* a two-species distribution model (see below), with independent data for *testing* model performance. The model is based upon 171 nominally high precision records (with two or more decimal places, from Dufresnes et al., 2020b). Model evaluation was done with the remaining records from that paper and the online data bases of the Societas Europaea Herpetologica (Sillero et al., 2014 and https://www.seh-herpetology.org/distribution-atlas), the Global Biodiversity Information Facility (available at https://www.gbif.org/species) and databases for Hungary and Croatia (Herpterkep, 2025 and https://biologer.hr/groups/16/species/126 and 127), two countries with otherwise low data coverage. Data for Hungary were kindly supplied by J. Vörös prior to formal release. Atlas grid cell data with both species were ignored in the analyses, as were records for Ukraine and Moldavia for which countries Corine land cover data are not available.

**Figure 1.**
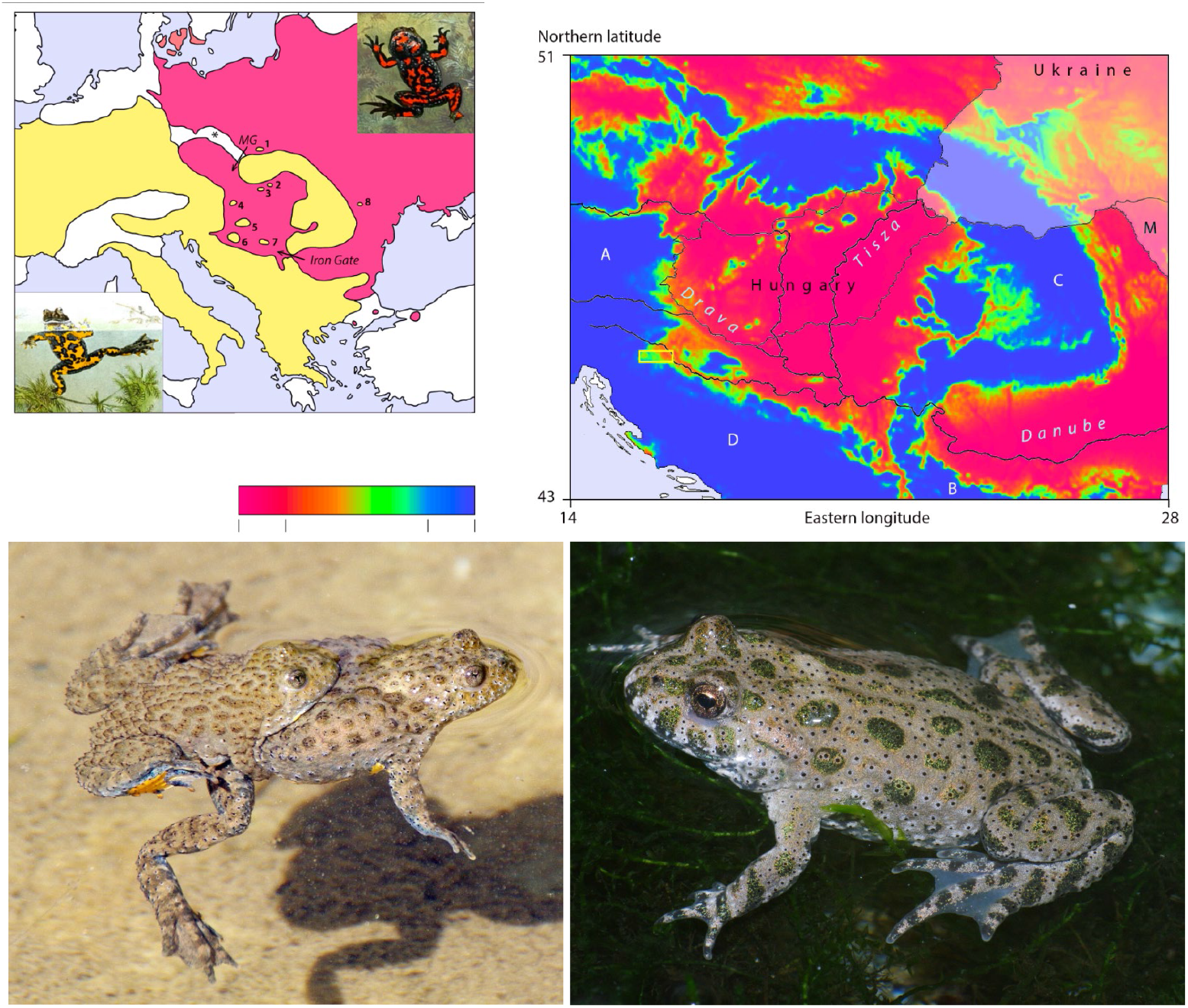
Panel top left – Distribution map for the red-bellied toad *Bombina bombina* (in red) and the yellow-bellied toad *B. variegata* (in yellow). Animal pictures are from Boulenger (1897) with *B. bombina* top right and *B. variegata* bottom left. Note the absence of either species in the Erz and Giant mountains (asterisk). Documented enclaves for *Bombina variegata* are at (1) the Kraków-Chrzanov ridge, (2) Aggtelek karst, (3) Mátra mountains, (4) Bákony forest, (5) Mecsek moutains, (6) Bilo Gora, (7) Fruška Gora and (8) Iaşi. Supporting data and recognition for the enclaves are by von Méhély (1905), Fuhn (1960), Michałowski (1961), Arntzen (1978) and Szymura (1993). Supporting data for the presence of *B. variegata* in northern Dobrogea, Romania (Mertens, 1928; Arntzen, 1978) have not become available and this occurrence was suppressed. Note that more recent and detailed data on the distribution of European *Bombina* species are available (e.g., Supplementary Information 2), but the production of an updated map is outside the scope of the paper. The arrow indicates the postglacial dispersal route of *Bombina bombina* across the Iron Gate from Dobrogea and the lower Danubian plains into the Pannonian plains (Vörös et al., 2006; Dufresnes et al., 2020b). An alternative but unsupported dispersal route follows the Moravian Gate (MG). Panel top right – Two-species distribution model (TSDM) for *B. bombina* and *B. variegata* in central Europe. Areas shown in red represent *B. bombina*, areas in blue represent *B. variegata* and ecological transition areas are shown in orange and green (0.2<*P*<0.8, see colour legend bottom left). The model is smoothened for presentation purposes. The main mountain systems are the Alps (A), Balkans (B), Carpathians (C) and Dinarides (Dinaric Alps, D) and the main rivers are the Danube, the Tisza and the Drava. For the Sava and Kupa rivers see Figure 2. Country borders of Hungary, Moldavia (M) and Ukraine are shown for reference (for other countries see Supplementary Information 2). Moldavia and Ukraine are white-shaded to indicate the absence of Corine land cover data. Bottom row left – amplexed pair of *B. variegata* from Greece. Bottom row right – *Bombina bombina* from the northeast of France (introduced population, Vacher et al., 2020). Photos by Serge Bogaerts.

**Figure 2.**
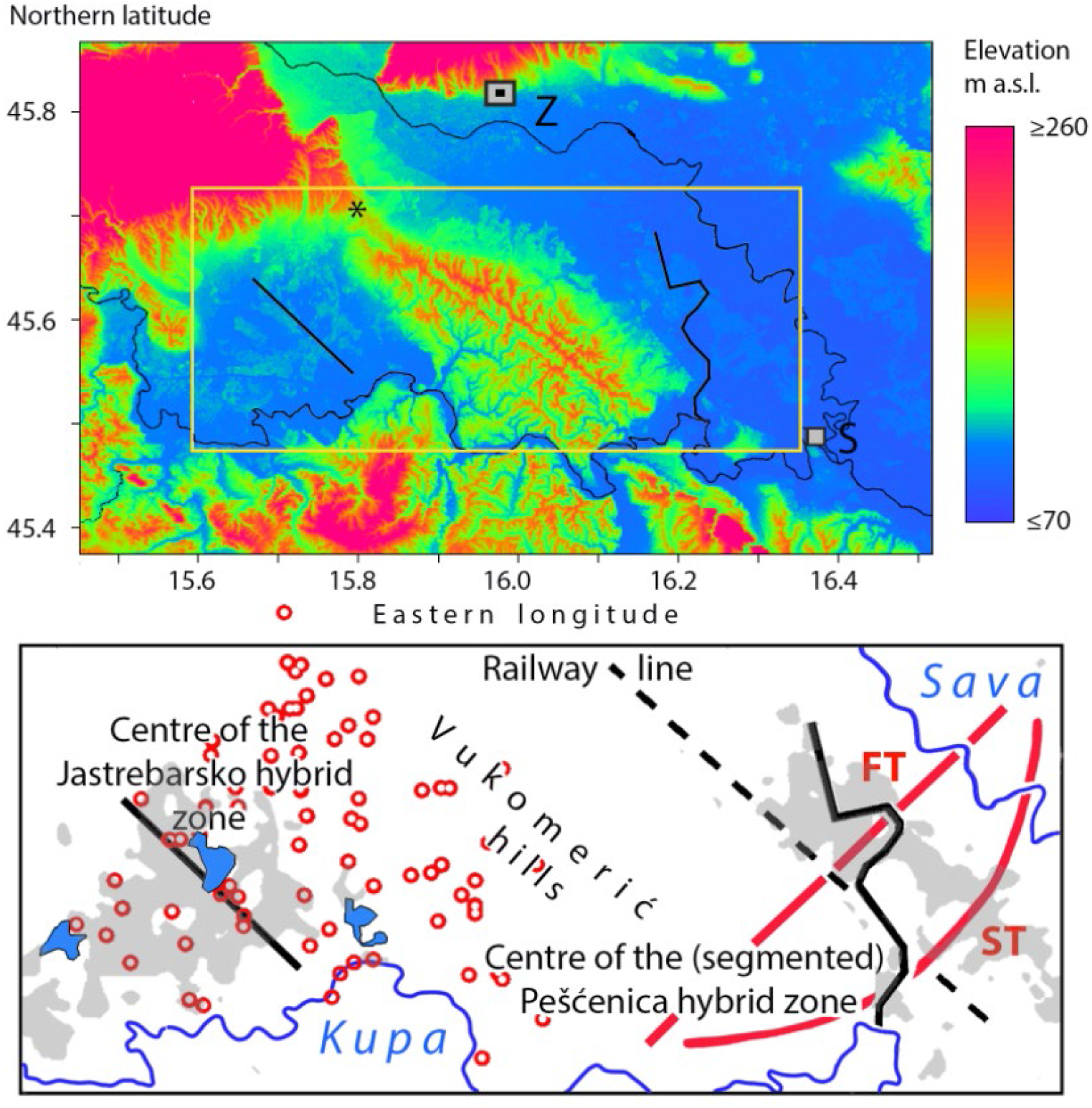
Study area of *Bombina* toads in Croatia south of Zagreb (Z) and west of Sisak (S). The top panel shows elevation from lower (in blue) to higher elevations (in red) with the adjoining Jastrebarsko and Pešćenica areas boxed. The thick black lines show the inferred centres of the *B. bombina – B. variegata* hybrid zones at either side of the Vukomerić hills and relative to the rivers Kupa and Sava and the Zagreb – Sisak railway line. The segmented hybrid zone at Pešćenica is after Atkinson (2001). The open red symbols in the lower panel are investigated *Bombina* populations in the Jastrebarsko area and the thick red lines summarize the forest transect (FT) and the southern transect (ST) at Pešćenica. Areas with lowland forest are shaded. Three lakes are shown by blue surfaces of which the central one is Crna Mlaka). Potential corridors that connect the pocket of *B. bombina* occurences at Jastrebarko with the main distribution in the Sava River lowland are over the saddle of the Vukomerić hills (asterisk) and the river Kupa.

Distribution data for central Croatia are as published (Atkinson, 2001) with species identifications based on a genetic hybrid index that runs from zero (pure *B. bombina*) to unity (pure *B. variegata*) with a 0.5 cut-off point (see also MacCallum et al., 1998). For the Jastrebarsko area species identification was based on ‘spot scores’ that describe the species’ differentiated ventral colouration patterns (Vörös et al., 2007 and references therein). Spot scores are concordant with results from genetic markers (Atkinson, 2001) and bimodally distributed with a classification cut-off point set at 0.45 (Figure 3).

**Figure 3.**
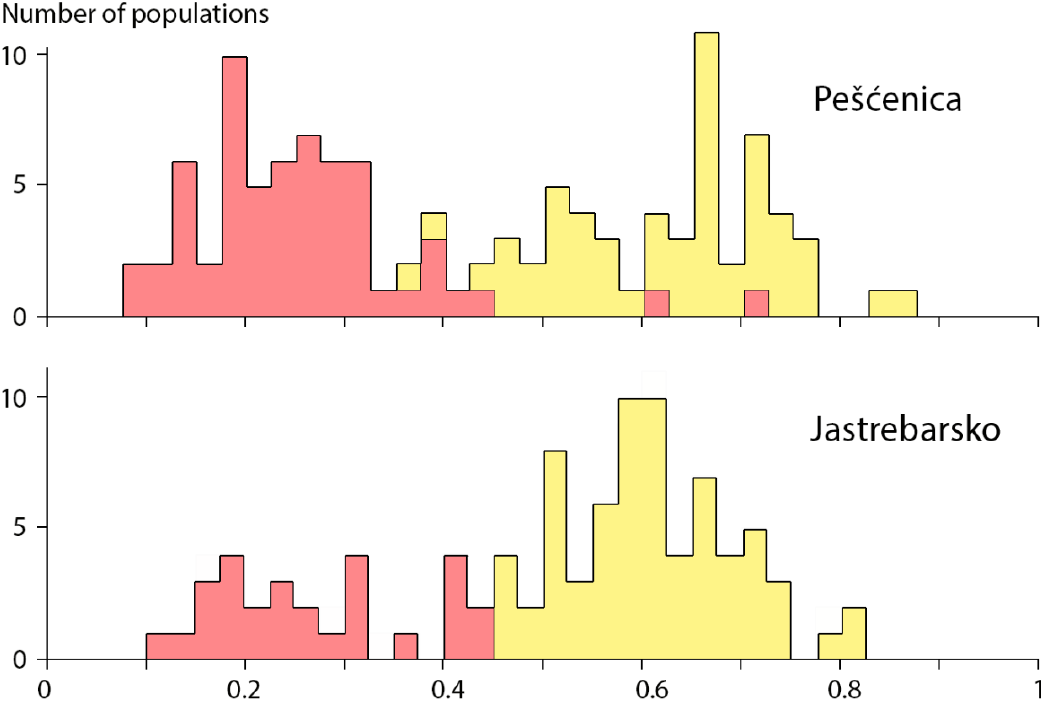
Histograms of spot scores of *Bombina* toads for the Pešćenica and Jastrebarsko areas in central Croatia. Data are taken from Atkinson (2001). The cut-off point is set at 0.45 based on genetic data from Pešćenica and then applied to Jastrebarsko, with *Bombina bombina* in red and *B. variegata* in yellow.

### Environmental data and two-species distribution modelling

Environmental data considered as candidate explanatory variables to the reciprocal distribution of *B. bombina* and *B. variegata* (and with that the position of the species’ hybrid zone) included 19 climate variables (bio01–bio19) extracted from the WorldClim global climate database v.2 (available at https://www.worldclim.org/data/index.html) (Fick and Hijmans, 2017). For elevation I used the Copernicus digital elevation model (European Space Agency, 2024 available at https://doi.org/10.5069/G9028PQB). ‘Hilliness’ is the standard deviation of elevation derived with a 9*9-pixelwide filter, covering 7.3 ha. Vegetation data were from the Corine land cover database of the European Environment Agency (available at https://land.copernicus.eu/pan-european/corine-land-cover, in particular https://doi.org/10.2909/71fc9d1b-479f-4da1-aa66-662a2fff2cf7) (Büttner et al., 2021). The nominal resolution of the data is 30 arc-seconds for climate, 30 m for elevation and 10 m for land cover. However, land cover data for Croatia were from 1998, corresponding to the period that the *Bombina* data were collected (available at https://doi.org/10.2909/c89324ef-7729-4477-9f1b-623f5f88eaa1). Following Arntzen et al. (2025), an *a priori* distinction was made between variables that operate at large spatial scales (i.e., the climate variables) versus elevation, hilliness and land cover that take effect at more local scales.

To identify and subsequently reduce collinearity among the environmental variables, a half-matrix of the pairwise absolute Spearman correlation coefficients was subjected to clustering using the unweighted pair group method with arithmetic mean clustering method in Primer 7 (Clarke and Gorley, 2015). Variables were retained using criteria of partial independence at |*r*|<0.75 and selected in alphanumerical order (Supplementary Information 1). Two-species distribution models (TSDM) in which the presence of one species is contrasted with the presence of the counterpart species, were derived with stepwise logistic regression analysis in SPSS v. 30 (IBM Corp., 2024). Parameter selection was in the forward stepwise mode under criteria of entry (P_in_=0.05) and removal of terms (P_out_=0.10) under the likelihood ratio criterion, while applying a weighing procedure that sets the number of records for species at par. The fit of the model to the underlying data was assessed by the area under the curve (AUC) statistic. Spatial models were analysed and visualized with ILWIS v.3.8.6 (ILWIS, 2019).

### Hybrid zone analysis

Classic equilibrium cline models, describing a sigmoid change in phenotypes or allele frequencies across hybrid zones (Szymura and Barton, 1991), were fitted with the R package HZAR (Derryberry et al., 2014) with details as described earlier (e.g., Arntzen et al., 2016). At Pešćenica two transects were studied from the lowlands into the hills as guided by a segmented *Bombina* hybrid zone reconstruction (Figure 2). The transects were projected through a lowland forest (the ‘forest transect’ with 40 investigated populations) and no forest adjoining the hills (the ‘southern transect’ with 27 populations). The species profile is made up of four diagnostic allozyme genetic markers. At Jastrebarsko, a single, wide transect was studied with spot score data for 78 populations (leaving out 19 populations with southwestern and southeastern positions that belong to an opposite stretch of the species contact zone, Supplementary information 3). Spatial reference is to the long straight stretch of the Zagreb-Sisak railway line that runs in parallel to the Vukomerić hills.

## Results

The TSDM for *Bombina* toads across central Europe is described by *P*=(1/(1+exp(0.0109*elevation+0.0437*hilliness-2.159*forest_cover+0.0128*bio12-12.494))) in which ‘bio12’ describes annual precipitation. The model fit is AUC=0.949±0.016. The position of the species contact zone corresponds to range borders reconstructed earlier, whereas the width of the ecological transition (0.2<*P*<0.8) varies widely (Figure 1). The fit of four sets of reference data to the model ranges from 0.939<AUC<0.962 (for a detailed account see Supplementary information 2).

Application of the spot score criterion to the Jastrebarsko area (where genetic data are unavailable) suggests that ca. a third of populations is at the *B. bombina* side of the species bimodality (Figure 3). Out of 12 environmental parameters available for selection (uncorrelated to the level at |r|=0.75, Supplementary Information 1) just elevation and forest cover made it into the model as *P*=(1/(1+exp(0.251*elevation-0.084*forest_cover-26.081))) with a model fit of AUC=0.902±0.035 (Figure 4). Testing the model with reference data was not possible for the lack of records for central Croatia (cf. Supplementary Information 2 panel F).

**Figure 4.**
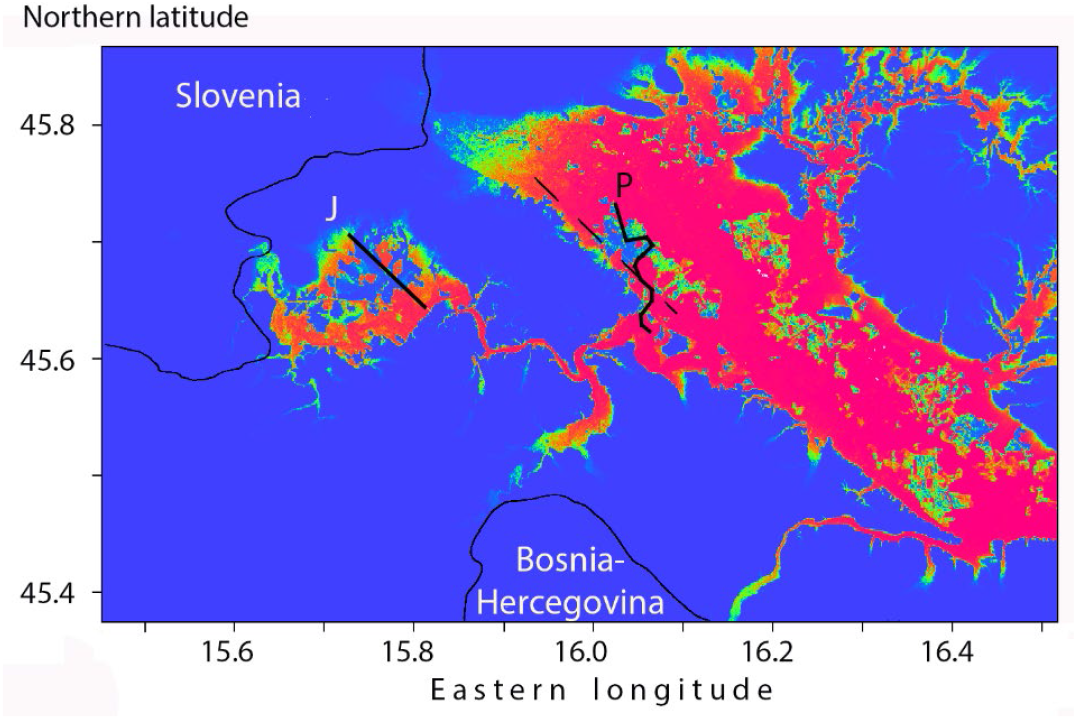
Two-species distribution model developed for *Bombina* species central Croatia. Areas shown in red represent *B. bombina*, areas in blue represent *B. variegata* and ecological transition areas are shown in orange and green (see colour legend in Figure 1). The centre of the hybrid zone at Jastrebarsko is shown by a solid straight line (J) and for Pešćenica by a segmented solid line (P). The railway line that is used as a spatial reference is shown by an interrupted line. The model supports the notion that the Jastrebarsko lowlands were colonized by *B. bombina* following the Kupa River valley in the south and not across the saddle of the Vukomerić Hills in the north.

At Jastrebarso the centre of the cline is positioned at a parallel distance of −8.8 km from the railway line at an altitude of ca. 125 m (Figures 4 and 5). A two-dimensional plot of spot scores at Jastrebarsko is near-equivalent to the linear hybrid zone representation (Supplementary Information 3 and 4). At Pešćenica the cline centres for two transects are positioned at either side of the railway line, set apart over 5.4 km and positioned at altitudes of ca. 111 m for the forest transect and ca. 105 m at the southern transect. Estimated hybrid zone widths range from 1.44-3.32 km (for details see Supplementary information 4).

**Figure 5.**
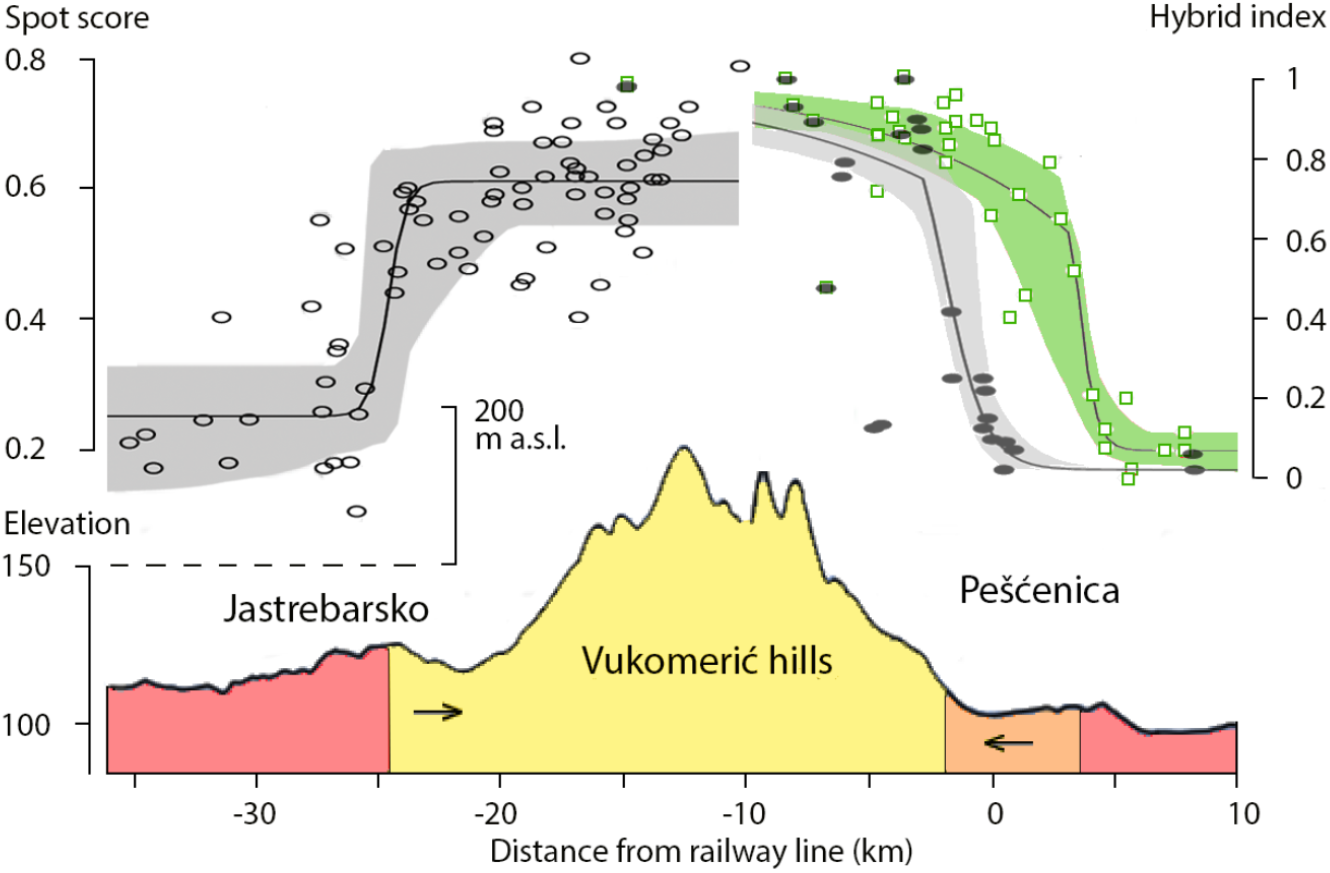
Distribution of *Bombina bombina* and *B. variegata* across the Vukomerić hills, combined for the Jastrobarsko and Pešćenica areas in central Croatia. Top panel – hybrid zones as reconstructed with HZAR software based on spot scores at Jastrebarsko and a genetic hybrid index at Pešćenica. Results for the forest transect (FT, in green) and southern transect (ST, in light grey) populations are shown by open square and solid ellipsoid symbols, respectively. The shaded area represents the 95% credibility interval of the estimate. Bottom panel – elevation profile. Areas are coloured yellow or red according to the majority presence of one species or the other (*Bombina bombina* in red and *B. variegata* in yellow). The orange part of the profile at Pešćenica is occupied by *B. variegata* across the forest transect and by *B. bombina* across the southern transect. The hypothesis eventually to be tested is that the hybrid zones would find a new position at the foothills if the local lowland forests were to become disconnected from the *B. variegata* range, or disappear altogether (see arrows).

## Discussion

### Species distribution modelling

Single-species distribution modelling is a popular research tool, also frequently applied to species for which the full ecological amplitude cannot be determined, such as when the species range is bounded by that of a closely related species. This then is unfortunate because, if species are affected to the extent that their ranges are interdependent, such a biotic interaction is better not ignored. It has, for example, been found that related salamander species can occur in a wider range of conditions than found at their parapatric contact (Werner et al., 2013). Similarly, for *B. variegata* it was observed that its ecological amplitude is wider in allopatry than in parapatry with *B. bombina* (von Méhély, 1905; Mertens, 1928) for the present day exemplified by remnant populations in the southwest of France (Berroneau et al., 2010) and in the Po Valley in Italy (Barbieri et al., 2004) and a thriving introduces population of *B. bombina* is known for northeast France (Vacher et al., 2020). Two-species distribution modelling highlights the ecological differences that come into play among competing taxa and neatly aligns with hybrid zone and species delimitation research (Arntzen, 2023a). Yet, modelling results away from the contact zone are possibly compromised and at the extreme a TSDM will project species where there are none, such as for *Bombina* toads in the Erz and Giant Mountains (Figure 1).

The ‘global’ TSDM for European *Bombina* includes the parameters elevation, hilliness, forestation and precipitation and is strongly supported by reference distribution data (Supplementary Information 2). The three locally operating parameters elevation, hilliness and forestation provide the model with substantial projected detail. The model is nevertheless frequently inaccurate such as, for example, at the Mecsek Mountains in southern Hungary where the realized *B. variegata* enclave is larger than projected (see panels A and F in Supplementary Information 2). This is probably because the *Bombina* hybrid zone is not positioned at one particular elevation contour. While the model sets the elevation separating the species at around 300 m, in the field it varies from ca. 115 m in central Croatia (present study) to ca. 330 m in western Ukraine (Yanchukov et al., 2006) and to ca. 450 m at the Hungarian-Slovakian border (Gollmann et al., 1988). In mosaic hybrid zone sections an elevation threshold is more difficult to determine, but for example at Apahida, Romania a hybrid belt is found at ca. 450 m (Vines et al., 2003). Another example of elevation operating at different thresholds is shown in the *Bufo bufo – B. spinosus* hybrid zone that is positioned a ca. 200 m in the west of France, at ca. 500 m at the Plateau Central and at ca. 1200 m in the Ligurian Alps (Arntzen et al., 2020). In these systems, the mutual species borders seem better described by the transition from flat to hilly than by elevation *per se* and global models are limited in the fit it can achieve. This flaw suggests that perhaps ‘hilliness’ is a better model predictor than elevation (Arntzen, 1996), but this is not obvious from the results because in a parallel study with Hungarian atlas data, hilliness yielded spatial extrapolations inferior to elevation (Arntzen et al., 2025).

For central Croatia only the parameters elevation and forestation make it into the *Bombina* TSDM. Visual inspection of the positioning of the hybrid zone at Pešćenica shows that the elevation contour is followed, except where extensive lowland forestation is connected to the *B. variegata* range (Bugter et al., 1997; MacCallum et al., 1998; Atkinson, 2001). A wider *B. variegata* range than predicted from the lowland to mountain transition is also found at Jastrebarsko (Figures 2, 4 and 5). The notion that forestation extends the position of the species hybrid zone into the lowland such as at Pešćenica – thus in favour of *B. variegata* – is here supported by analyses at large (central Europe) and small spatial scale (Jastrebarsko). The importance of forestation to the presence of *B. variegata* is further supported by a TSDM for *Bombina* species from the Hungarian atlas data (Arntzen et al., 2025). The transects in central Croatia yield hybrid zones that at 1.4-3.2 km are narrower than elsewhere (average width 7.0 km, range 2.1-11.4, data compiled by Dufresnes et al., 2020b) and it is probably not coincidental that the other most-narrow hybrid zone at Strych, Ukraine with a width of 2.1 km also yields an intricate and convoluted parapatric species contour (Yanchukov et al., 2006: Figure 1C).

### From Holocene to the present

Consensus exists that during the most recent Pleistocene glaciation at 20 Kyr BP refugia were in Dobrogea and at the Black Sea coast for *B. bombina* and in the southern Carpathians and the Balkan and Apennine Peninsulas for *B. variegata* (Arntzen, 1978; Dufresnes et al., 2020b). A Pannonian refugium for *B. bombina* is contradicted by the occurrence of enclaves of *B. variegata* in this area, suggesting that the latter species was there first (Vörös et al., 2016). The Holocene colonization of Pannonia by *B. bombina* is thought to have taken place from the southeast, following the Lower Danube Corridor through the Iron Gate (Arntzen, 1978; Magyari et al., 2010). While Dufresnes et al. (2020b) support this scenario, they assume that studied sections of the *B. bombina* - *B. variegata* hybrid zone formed at their current position shortly upon post-glacial climate amelioration. I take issue with this detail because it ignores the buffering effect of forestation on the *B. bombina* range expansion. The research question here addressed parallels the debate on the biogeographic origin of the species-rich steppe grasslands in central Europe, to which the alternative hypotheses are long-term species persistence in situ versus immigration from the south-east, either after the last glacial maximum (LGM) or after the Neolithic landscape deforestation (Divíšek et al., 2022).

Most of Pannonia and the Danube Plains was forested at 6 Kyr BP (Bohn, 2003; Molnár et al., 2018; Feurdean et al., 2021). The prime *B. bombina* habitats of wetland areas and open steppes were located mostly adjacent to the Danube and Tisza rivers (Figure 6). Given the relatively strong position of *B. variegata* in lowland forests observed in the present study, these forests may have been strongholds that remained occupied by *B. variegata* for longer than the surrounding deforested areas. An extensive patchwork of *B. variegata* occurrences may have existed over the Pannonian and Danubian Plains of which forested patches eventually dissolved and elevation enclaves persisted. However, ambiguity exists on tempo and modo of the deforestation process. Anthropogenic influences started possibly as early as 3100 yr BP (Magyari et al., 2010) yet woodland cover was high until the Middle Ages (Sundseth, 2009) and the absence of *B. variegata* in the Pannonia lowland may have been relatively recent. By the late 18^th^ century forestation in Pannonian lowland was down to ca. 10% cover, further down to ca. 3% in 1942, with a recent upsurge of secondary forest at the expense of semi-natural forest (Biró et al., 2022). The relative recency of the deforestation process and the subsequent latter-day advance of *B. bombina* over *B. variegata* offers opportunities for molecular genetic research (see below).

**Figure 6.**
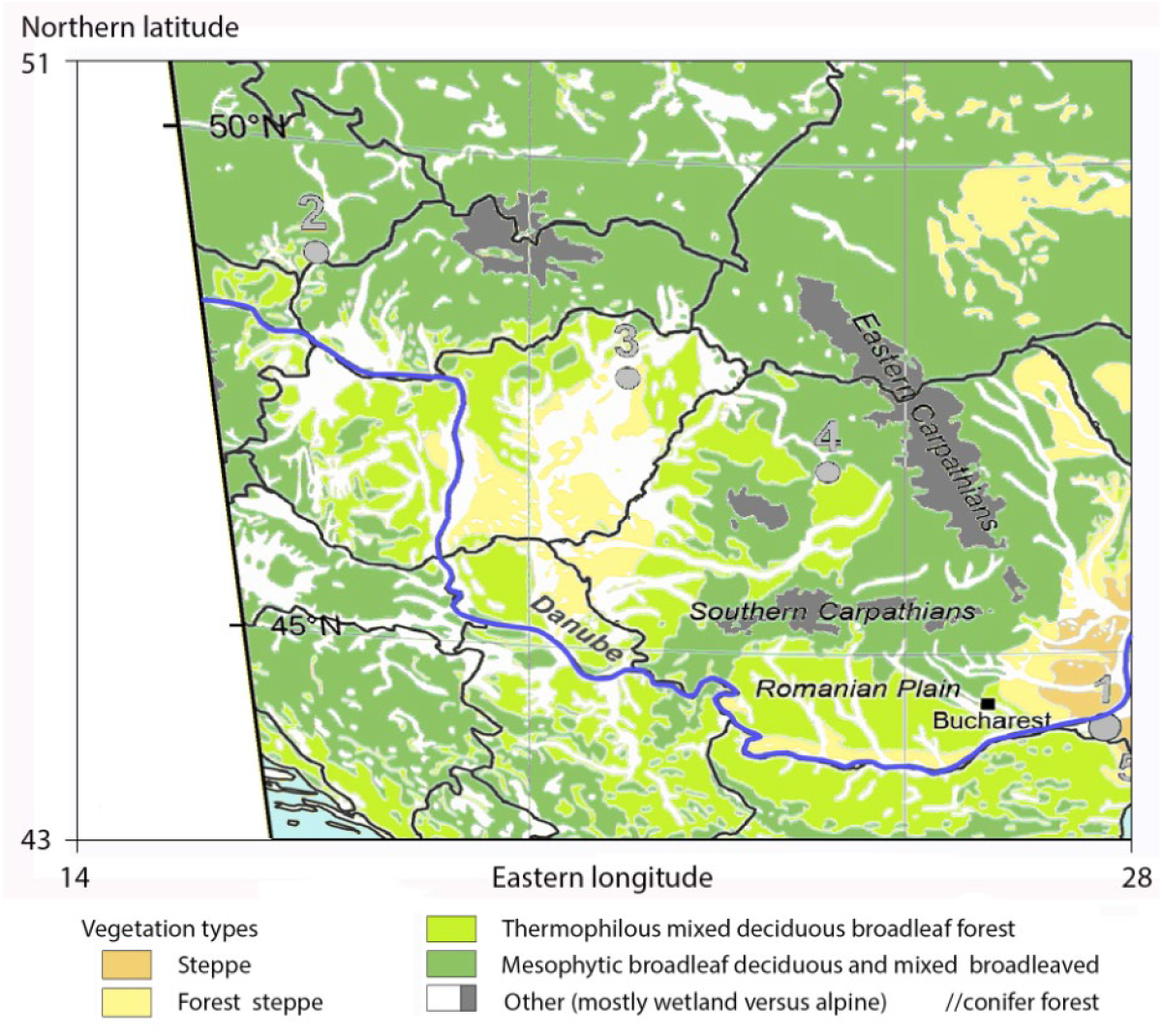
Potential natural vegetation cover showing the extent of wetlands, steppes and forestation in central Europe at 6000 yr BP (after Bohn, 2003 and Feurdean et al., 2021). *Bombina bombina* prime habitat is unforested lowland and steppes wetland, as here shown in orange and white. Forested steppes and other forests (in yellow and green) at that time widespread over the current range of *B. bombina* (cf. Figure 1) were possibly inhabited by *B. variegata*.

The Jastrebarsko area forms a miniature representation of the *Bombina* area of range overlap across central Europe, with a potential corridor in the north (the saddle at the Vukomerić hills versus the Moravian Gate that interconnects Poland and the Czech Republic) and a river that forms the actual corridor (Kupa River valley versus the Danube at the Iron Gate) (Figures 1 and 2). However, at Jastrebarsko the presence of *B. bombina* is limited to the Kupa River region whereas the Pannonian Plain is almost entirely taken. The hybrid zone at Jastrebarsko runs through the centre of the lowland forest at ca. 5 km from the lowland-mountain transition, so that areas close to the Kulpa River have *B. bombina* and areas adjacent to the Vukomerić hills have *B. variegata* (Figure 5). A complicating factor in this semi-isolated lowland pocket *–* for which I propose the term ‘peninsular enclave’ or ‘pen-enclave’ *–* is the presence of three fish pond complexes (Figure 2) that are (or were) *B. bombina* habitat and are avoided by *B. variegata* (von Méhély, 1905; Dolgener et al., 2012).

*Bombina bombina* and *B. variegata* are closely related species that engage in extensive hybridization. The species are also deeply differentiated across a wide variety of life history traits (see Szymura, 1993: Table 1). A fundamental difference between the species is the aquatic versus more terrestrial lifestyle that is reflected in typical breeding habitats found at opposite ends of the large-and-permanent (for *B. bombina*) to small-and-ephemeral continuum (for *B. variegata*) (Supplementary Information 6). The emphasis on the breeding habitat as a local factor that affects the species’ interaction and local distribution (MacCallum et al., 1998; Vines et al., 2003; Yanchukov et al., 2006) is informative but temporally biased because most puddles are ruts and car-track from forest management whereas large, stable ponds are frequently purposefully made. Such anthropogenic influences may have assisted the persistence of *B. variegata* in forests and of *B. bombina* in lowlands, therewith strengthening the species’ ecological segregation.

**Table 1.**
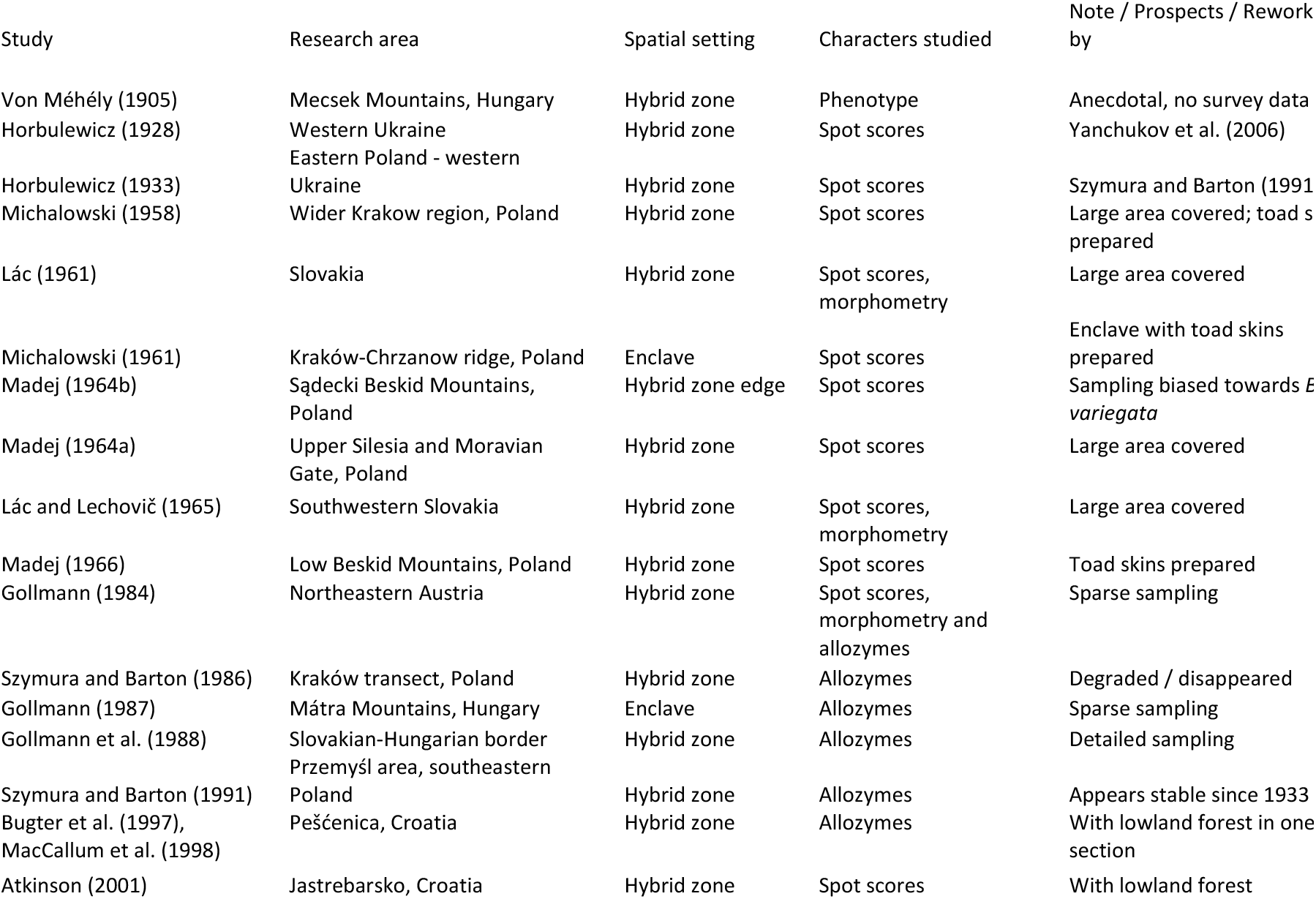
Overview of historical studies with fieldwork before 2000 on *Bombina bombina* and *B. variegata* hybrid zones, considered for resurveying and additional research.

The combined effect of elevation and forestation on species dynamics as here shown for *B. bombina – B. variegata* is known from other systems. European amphibian species pairs in which a lowland species supersedes a closely related species from the hills, mountains and forests include *Triturus cristatus* and *T. pygmaeus* sandwiching *T. marmoratus* (Arntzen and Wallis, 1991; Espregueiro Themudo and Arntzen, 2007; Arntzen and Espregueiro Themudo, 2008; Arntzen, 2023ab) and *Bufo bufo* displacing *B. spinosus* (Arntzen, 2019; Arntzen et al., 2020). Another candidate pair for the study of species replacement by a hybrid zone moving up an elevation-forestation gradient would be *Lissotriton vulgaris* - *L. montandoni* (Antunes et al., 2023ab). This raises the question if the reverse situation of a mountain/terrestrial species replacing a lowland/aquatic species is just more difficult to document or that these cases are rare.

### Prospects for research

If the *Bombina* hybrid zone is set at an elevation gradient, than the current situation possibly formed shortly after post glacial climate warming (Arntzen, 1978; Dufresnes et al., 2020b). However, as suggested in the present paper, forestation prevented or tempered the *B. bombina* advance. The process is recent (a couple of centuries) and may be ongoing. A hybrid zone not at equilibrium but moving opens research possibilities. Three lines of argument have successfully been employed to the study of species replacement by moving hybrid zones, namely (i) historical data, (ii) spatial patterns of relict populations and (iii) introgression of genetic markers into the advancing taxon (Arntzen and Wallis, 1991; Buggs, 2007; Wielstra, 2019). For the *Bombina* research program this may turn out as follows.

#### Historical data

The resurveying of documented *Bombina* distributions (Horbulewicz, 1928, 1933) was carried out in southeastern Poland and western Ukraine (Szymura and Barton, 1986; Yanchukov et al., 2006) (Table 1). This yielded neutral results of no change, presumably because hybrid zones locally stabilized against an elevation profile. Among another seven studies with spot score data (1958-1966) the Polish ones stand out for a wide coverage. An especially promising setting for resurveying is the *B. variegata* enclave at the Kraków-Chrzanow-ridge (Michałowski, 1961; see Figure 1 and Supplementary Information 5), the more so if the toad skins that Michałowski prepared would still be available, as a reference and to genetically corroborate his spot score classification. Among studies from the allozyme era (1984-1998) the one at the Slovakian-Hungarian border stands out for sampling detail (Gollmann et al., 1988). Especially promising venues are lowland forests close to or connected with the continuous *B. variegata* range such as at Pešćenica and Jastrebarsko, although it remains to be seen if not the time span of three decades or less is not too short for movement of the hybrid zone to be detectable. Elsewhere, either sampling was sparse or transect habitats have become derelict (Table 1). Mosaic hybrid zones such as at Pahadia, Romania (Vines et al., 2003) are promising because of possibly fluctuating environmental conditions, but require detailed sampling to cope with the two dimensions.

#### Spatial patterns

Both *Bombina* species have ranges with isolated populations at the fringes of their ranges, such as for *B. variegata* across France (Lescure et al., 2011; Vacher et al., 2020) and for *B. bombina* in European and Asiatic Turkey (Figure 1) suggesting range regressions. Other isolated *B. variegata* occurrences are enclaves where the species is surrounded and thus isolated by *B. bombina*. These are an indication of the species’ wider past distribution and the advance of its counterpart *B. bombina* (Arntzen, 1978). The most promising areas for discovering enclaves are elevated, forested and data-deficient areas in northern Rumania, western Ukraine and Moldavia. So far, an association with *B. variegata* was found in the floodplain beech forests near the Dniester River in Ukraine (possibly as far east as the Medobory hills) and the Buciumeni forest in Rumania (Supplementary Information 2 panel F) but more data are needed to resolve more *B. variegata* enclaves, if they exist.

#### Genetic footprints

Species replacement may also transpire from ‘genetic footprints’ left behind by the regressing counterpart (Scribner and Avis, 1994; Currat et al., 2008). Dufresnes et al. (2020b) point to ‘genetic leakage’ in the *Bombina* system but reported no asymmetries that would be indicative of spatio-temporal directionality. A candidate area for research is the North Hungarian Mountain Range that is seemingly suitable for *B. variegata*, but where the species is present in some (Bakony, Mátra) and appears absent in other mountains (Bükk and Börszöny) (Arntzen et al., 2025). These areas may represent different stages in the process of species replacement and it is just a matter of opinion if the situation at Mátra (with possibly no pure *B. variegata*, Gollmann, 1987) is best described as either a deep genetic footprint or an eroded enclave. Other candidate research areas are persisting semi-natural lowland forests and areas where these disappeared recently, as can be traced in detail starting with the First Military Survey, that took place 1782–1785 (Hungarian Military History Museum, Budapest, in Biró et al., 2018, 2022). Under the prevalent scenario of an advancing *B. bombina* and a regressing *B. variegata* it is advisable that future molecular studies should include populations from the *B. bombina* refugium areas at the western and northern Black Sea coast, to confirm that perceived *B. variegata* genetic traces are not actually *B. bombina* ancient polymorphisms persisting through incomplete lineage sorting. Future genetic work on *B. variegata* enclaves may shed some light on the pattern and process of species replacement though the timing and the size and environmental features of the enclave will both affect genetic introgression in ways that may be hard to disentangle.

## Supporting information

Supplementary Informatio

## Conflicts of Interest

The author declares no conflicts of interest.

## Funding information

The start-up of this project was made possible by the Small Research Grant ‘Analysis of the distribution of the toads *Bombina bombina* and *B. variegata* across a hybrid zone: predictions from remotely sensed data’ (GR9/01719) of the UK Natural Environment Research Council, allocated to JWA and N. H. Barton.

## Data Availability Statement

This article does not contain new data. The data that are analysed are publicly available at: http://hdl.handle.net/1842/13662, https://biologer.hr/groups/16/species/126 and 127, https://doi.org/10.2909/71fc9d1b-479f-4da1-aa66-662a2fff2cf7, https://doi.org/10.2909/c89324ef-7729-4477-9f1b-623f5f88eaa1, https://land.copernicus.eu/pan-european/corine-land-cover https://www.gbif.org/species, https://www.seh-herpetology.org/distribution-atlas and https://www.worldclim.org/data/index.html, as also referenced in the main text.

## Supporting Information

Additional supporting information can be found online in the Supporting Information section.

## Notes

### Competing Interest Statement

The authors have declared no competing interest.

