## Supplementary Informatio for "Long term hybrid zone dynamics in red- and yellow-bellied toads estimated from environmental data at allopatric and parapatric scales"

**Supplementary information 1**

Clustering of environmental data with the unweighted pair group method with arithmetic mean on basis of the absolute value of Spearman’s correlation coefficient. Variables selected for two-species distribution modelling are indicated by a square symbol.

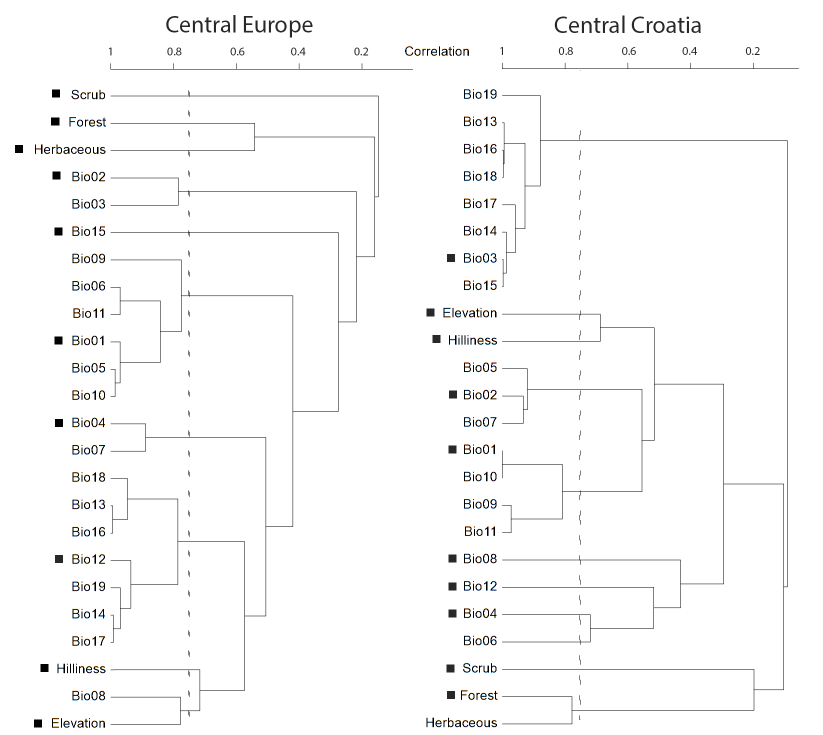

**Supplementary information 2**

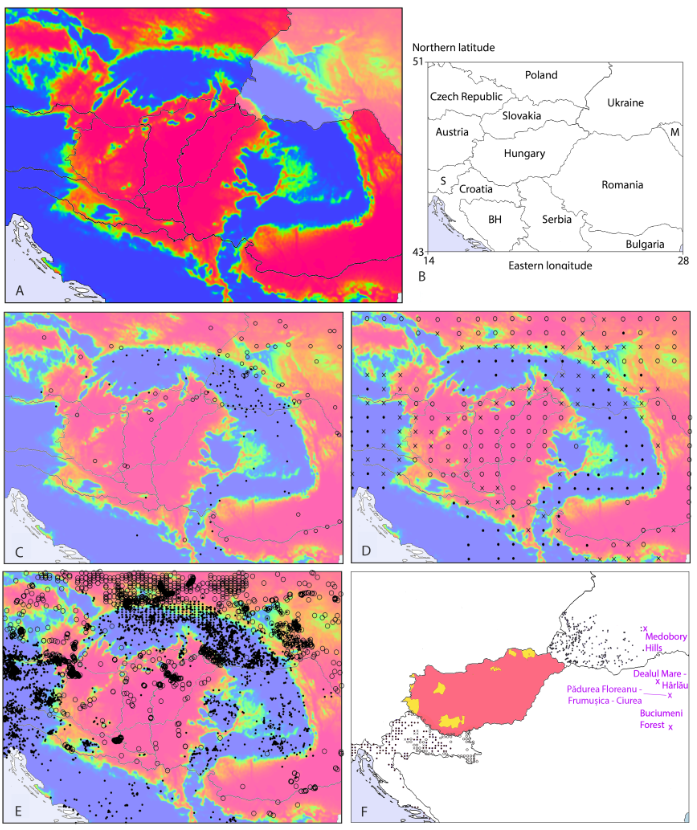

A – Two-species distribution model (TSDM) for *Bombina bombina* and *B. variegata* in central Europe as in Figure 1. Areas shown in red represent *B. bombina*, areas in blue represent *B. variegata* and ecological transition areas are shown in orange and green (0.2<*P*<0.8). The colour legend is as in Figure 1. The model is smoothened for presentation purposes. Moldavia and Ukraine are white-shaded to indicate the absence of Corine land cover data. B – Countries that together cover the area of *Bombina bombina - B. variagata* range overlap in central Europe. Abbreviations are: BH – Bosnia-Hercegovina, M – Moldavia, S – Slovenia. Panels C, D, E and F show the records used for testing the model as obtained from published data, with *B. bombina* shown by open round symbols, *B. variegata* by smaller, solid round symbols whereas both species are shown crosses (only in panels with grid cell data). Panel C – low resolution data from Dufresnes et al. (2020) with model fit AUC=0.939±0.019. Panel D – SEH atlas (Sillero et al., 2014) with model fit AUC=0.963. Panel E – GBIF data () with model fit AUC=0.949. Panel F – atlas data for Croatia (<https://biologer.hr>) with model fit AUC=0.963, the continuous distribution map of both species in Hungary as represented by Dirichlet cells with a model fit 0.954<AUC<0.983 (Arntzen et al., 2025) and records on *B. variegata* for Ukraine (Pisanets et al., 2005). Outermost eastern localities of B. variegata are the Medobvory Hills in Ukraine (S. Litvinchuk, pers. comm.), the site Dealul Mare - Harlau (see <https://eunis.eea.europa.eu/sites/ROSCI0076>), and the Buciumeni forest in Rumania (Pop et al., 2019) and possibly Pădurea Floreanu - Frumușica - Ciurea

**Supplementary information 3**

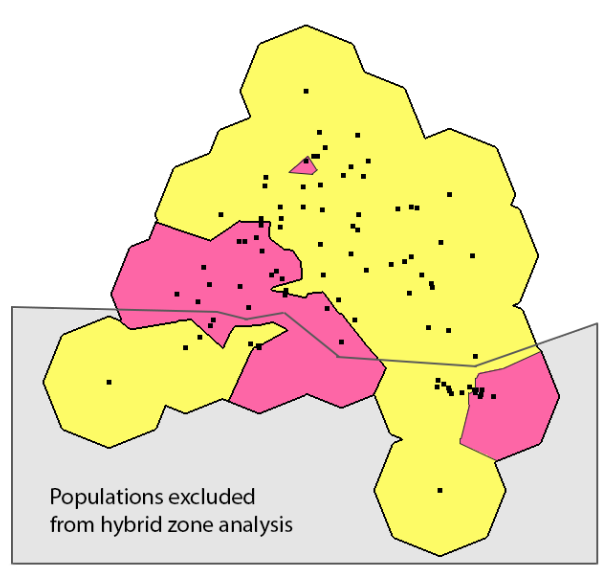

Two-dimensional plot of spot scores for the Jastrebarsko area for localities with *Bombina bombina* (in red, populations with spot score <0.45) and *B. variegata* (in yellow with spot scores ≥0.45). Note that populations to the south are excluded from analysis because they represent different sections of the species hybrid zone.

**Supplementary information 4**

Parameter estimates for the *Bombina bombina – B. variegata* geographical clines shown in Figure 5. Position is the centre of the contact zone relative to the Zagreb-Sisak railway line (see Figure 2) and cline width (w) is calculated as 1/maximum slope. Confidence intervals are two log-likelihood unit support limits. Tail fitting involved adjustments at the left side of the cline by shape parameters δ and τ. P_min_ and P_max_ are the character states at either end of the transect that are either fitted (‘opt’ models) or empirical values (the ‘fix’ model).

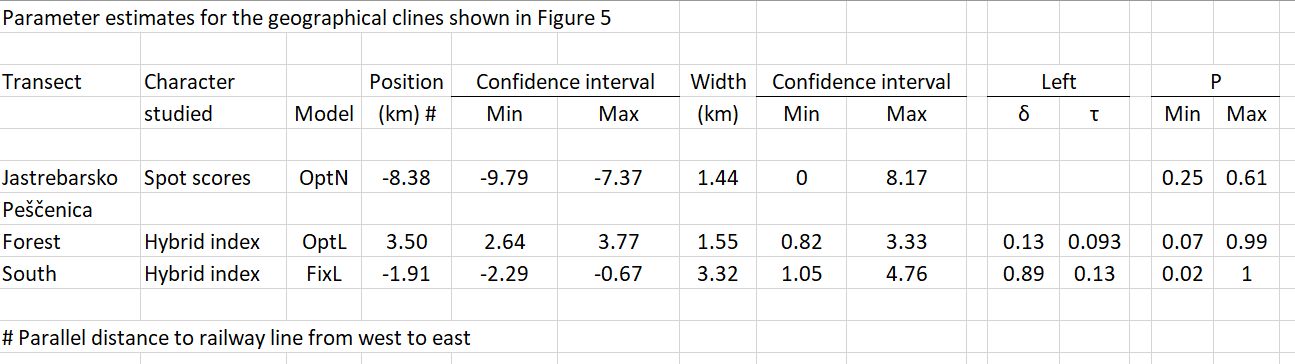

**Supplementary information 5**

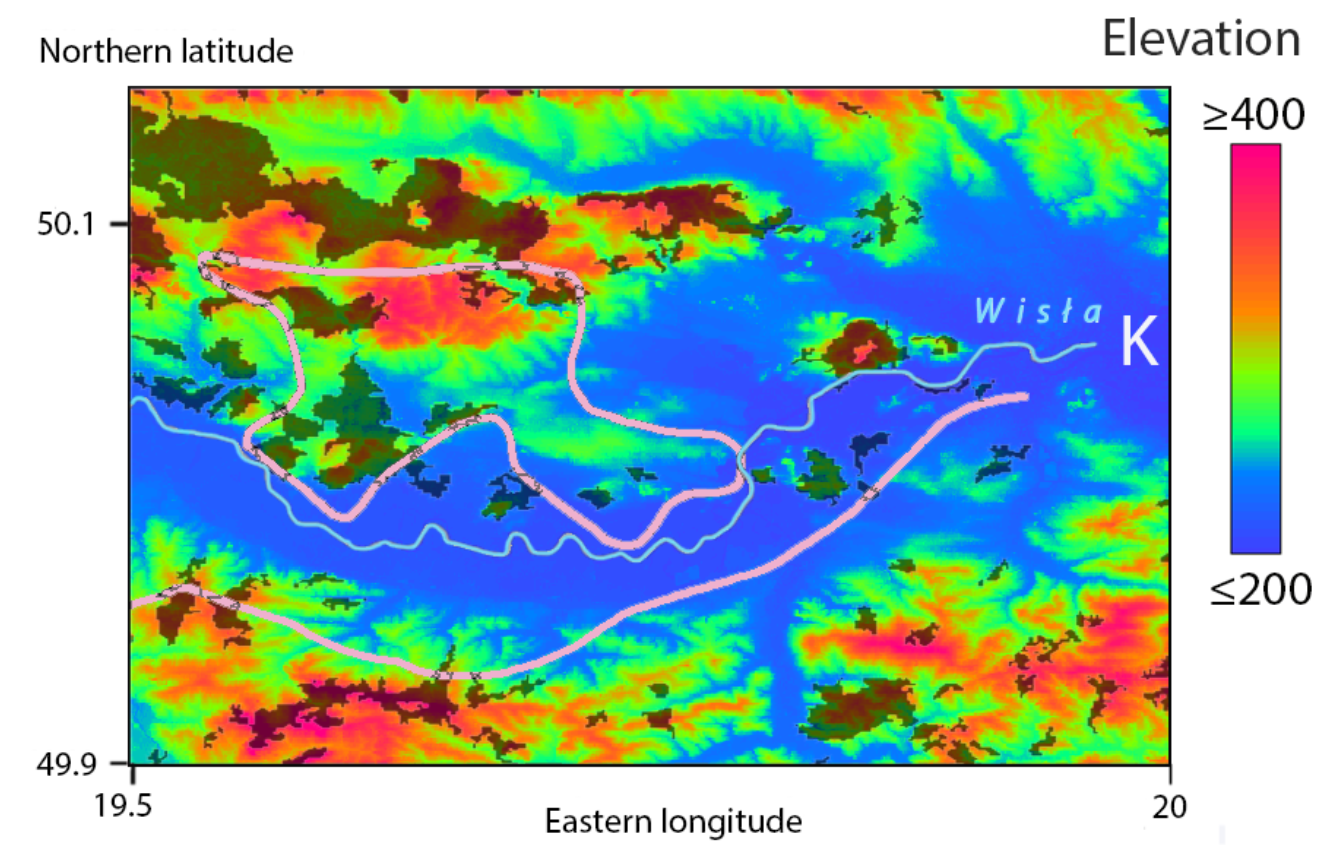

The distribution of *Bombina bombina* and *B. variegata* over the northern Carpathians and the Kraków-Chrzanów ridge according to Michałowski (1958), plotted over an elevation map (see colour legend). The fat pink lines show the northern edge the *B. variegata* range as well as an enclave at the ridge, north of the river Wisła. Forestation is grey shaded. K - Kraków.

**Supplementary information 6**

Breeding sites for European *Bombina* toads. Left – a medium sized, steady pond in a lowland gravel pit near Illmitz, Burgenland, Austria, breeding site for *B. bombina* (see Lörcher, 1969). Right – temporary car-track puddles in the Bakony forest, Hungary, breeding site for *B. variegata*.

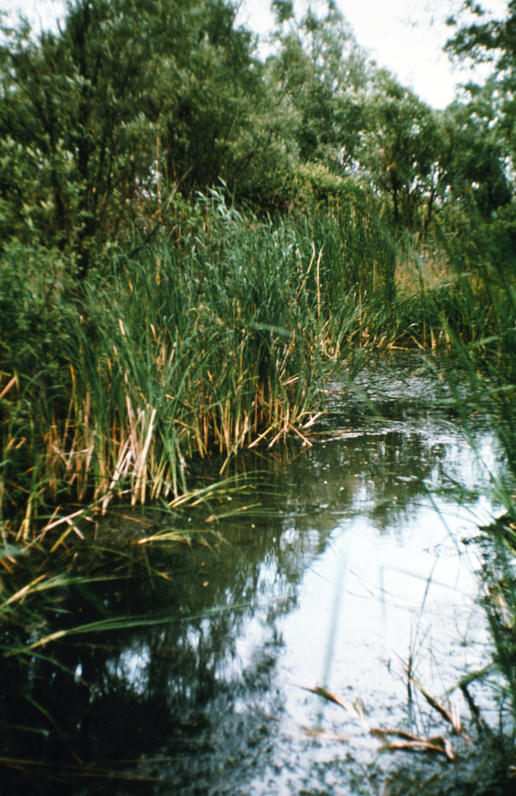

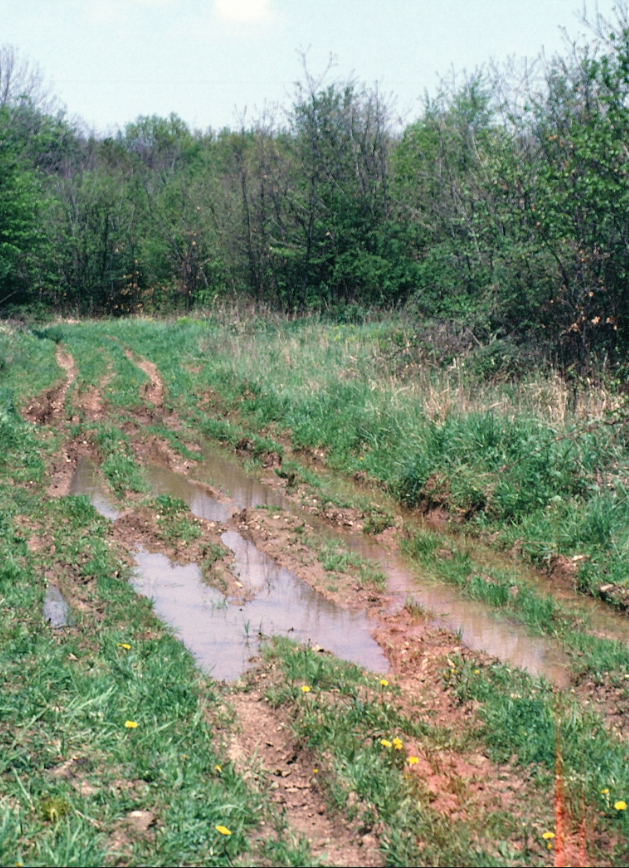
